# Breakthrough in Functional Muscle Regeneration: Biomimetic Collagen Scaffold Restores Abdominal Wall Defects in a VML-rabbit Model

**DOI:** 10.1101/2025.06.07.657281

**Authors:** Xiaoming He, Zhaohui Luo, Shenghua He

## Abstract

Functional regeneration of volumetric muscle loss (VML) remains a significant challenge in the field of regenerative medicine.

We have developed a novel biomimetic scaffold (0.4 mm thick hernia patch) derived from soluble collagen. This biomimetic scaffold exhibits exceptional strength, low immunogenicity, and structurally mimics the natural muscle fiber arrangement. The scaffold is utilized to repair a VML rabbit model (30 × 30 mm defect in abdominal wall) using nylon sutures for connection. It was observed that the VML site was gradually covered by regenerated tissue, which consisted of 2-3 mm thick functional muscle. By 32 weeks post-surgery, the newly formed muscle tissue had covered the majority of the VML area, as evidenced by morphological observations and histological evaluations. Approximately 24 weeks after surgery, the scaffold underwent complete degradation. This degradation exhibited a strong correlation with muscle regeneration. Furthermore, it was observed that the nylon sutures gradually migrated towards the VML center as muscle regeneration progressed, and nylon sutures exhibited an adverse impact on muscle regeneration..

The study did not employ exogenous cells or growth factors. However, the collagen scaffold effectively stimulated endogenous muscle regeneration. In contrast to the previous limited partial recovery or fibrotic scar healing, the collagen scaffold successfully induced genuine structural and functional muscle regeneration.

The 0.4 mm thick scaffold (hernia patch) exhibits a different structure from the 2-3 mm thick natural abdominal muscle wall. This suggests that a simple scaffold can regenerate functional tissue with a complex structure.

This research represents the first successful functional regeneration of VML in clinically relevant animal models through the utilization of artificial materials. This technology possesses transformative potential in the treatment of muscle defects arising from trauma, tumor resection, hernia, or congenital anomalies. Furthermore, it holds substantial medical potentials in addressing neuromuscular diseases, such as Duchenne muscular dystrophy (DMD), amyotrophic lateral sclerosis (ALS), and spinal muscular atrophy (SMA), by collaborating with advanced technologies including CRISPR-Cas9, adeno-associated virus (AAV) gene therapy, and stem cell technology.

## Introduction

Volumetric muscle defect (VML) is a serious and usually irreversible disease, which is defined as the loss of skeletal muscle tissue that exceeds the body’s innate regenerative ability and cannot rely on itself to achieve complete regeneration of structure and function [Testa et al., 2021]. VML is usually caused by trauma, tumor resection or surgical complications, such as abdominal wall hernia. VML sites not only have structural deformity and chronic pain, but also have serious loss of function. The patient’s quality of life declines, and often need to be operated again [Petro et al., 2015; Greising et al., 2019].

The need to treat VML and effectively regenerate missing muscles is both urgent and growing. Worldwide, there are more than 6.8 million surgeries involving muscle resection or large fascia injury every year [Testa et al., 2021]. In the United States alone, there are more than 350,000 abdominal wall reconstruction surgeries every year, and this number is expected to increase further due to the increase in obesity, caesarean section and cancer resection rates [Sanders and Kingsnorth, 2012]. It is expected that by 2028, the global market for soft tissue repair materials will exceed $9.3 billion, of which muscle regeneration is one of the fastest growing submarkets [Kulwatno et al., 2023].

Despite the pressing need for VML treatment, the current clinical treatment modalities encounter substantial limitations. Autologous muscle tissue transplantation is the main means of treating VML, however, autologous muscle tissue transplantation also presents limitations. For instance, the arteries, veins, and nerves of the donor’s site are severed, leading to compromised biomechanics and functional impairment and defects in the donor area. However, the treatment outcome is unsatisfactory, and normal anatomy is also lost in the repair area [Grasman et al., 2015]. Furthermore, its own muscle tissue is limited, which is insufficient to meet the requirements of VML transplantation. Consequently, researchers attempted to utilize alternative materials, such as decellularized tissue, including pig’s small intestine submucosa (SIS), pericardium or pericardial tissue, urinary bladder matrix (UBM), blood vessels, valves, broad fascia, and dermis. These materials are employed to repair VML [Butler and Prieto, 2004; FitzGerald and Kumar, 2014; Brown and Badylak, 2014; Corona et al., 2014; Overbeck et al., 2020; Garvey et al., 2017; Koscielny et al., 2018; Harris et al.,2021; Olavarria et al., 2021; Kish et al., 2005]. However, the efficacy of these methods is limited and lacks the ability to genuinely regenerate functional skeletal muscles. Decellularized matrix contains various antigenic components, and even completely decellularized collagen framework contains antigenic telopeptides, which may influence the migration, growth, and differentiation of satellite cells (muscular stem cells). Decellularized heterogeneous muscle tissue fails to successfully regenerate muscle tissue [Badylak et al., 2009; Turner and Badylak, 2013]. Even autologous devitalized muscle tissue has demonstrated insufficient efficacy in regenerating muscle tissue as a treatment for VML [Garg et al., 2014].

Synthetic polymer materials have gained significant attention in the field of muscle regeneration, particularly in the realm of hernia repair involving the abdominal wall muscles. Among the commonly utilized materials are non-degradable polypropylene (PP), expanded polytetrafluoroethylene (ePTFE), and polyester (PE). These materials are favored for their stability, absence of disease transmission risks, high toughness, moderate flexibility, compatibility, and low cost. However, these materials cannot restore the original muscle structure and function, furthermore, certain complications may arise, including a high recurrence rate, foreign body reactions, scar tissue formation, adhesions, intestinal obstruction, erosion, chronic pain, chronic infection, intestinal fistula, and other adverse effects [Grasman et al., 2015; Aydemir et al., 2019; Lee et al., 2020; Usher et al., 1959; Borrazzo and Prieto, 2004; Matthews et al., 2003; Fuziy et al., 2019]. On the other hand, degradable materials such as polylactic acid have been abandoned due to the high incidence of fistula, but some researchers are trying to combine them with other materials [Xv et al., 2025]. In addition, compatible synthetic materials, hydrogel, stem cell transplantation, addition of antibiotic and growth factors, precise microstructure control and bioprinting method are also applied in muscle tissue engineering. These techniques have a certain effect on small animal models or partial damage repair, but they have not successfully reconstructed functional and aligned muscle fibers in VML site, especially in large animal models [Wang et al.,2020; Hwangbo et al., 2021; Blatnik et al., 2017; Yabanoğlu et al., 2015; Wang et al.,2020; Passipieri and Christ, 2016; Wang and Yang 2020].

The primary challenges confronting the field of muscle bioengineering include insufficient mechanical strength of the scaffold, absence of muscular structural alignment, immune response, and limited in vivo vascularization [Testa et al., 2021; Greising et al., 2019; Sanders and Kingsnorth, 2012; Kulwatno et al., 2023; Gilbert-Honick and Grayson, 2020].

Collagen, a crucial protein component of muscle structure, undergoes antigenic telopeptide removal to produce soluble collagen. This soluble collagen possesses exceptionally low antigenicity, high affinity, and abundant ‘nutritional value’, making it a globally recognized excellent medical material [Sowbhagya et al., 2024]. However, its limited strength, strong hydrophilicity, susceptibility to expansion, and rapid degradation in vivo restrict its further application.

In response to the aforementioned challenges, we have developed a novel biomimetic scaffold (hernia patch) based on soluble collagen, mimicking the structure of muscle fibers. This scaffold exhibits high affinity, low antigenicity, exceptional strength, and a slow degradation rate in vivo. The hernia patch was utilized to repair the VML in the abdominal wall of a rabbit model. The results demonstrate functional muscle regeneration in the VML site without the use of exogenous cells or growth factors.

This report presents the inaugural functional muscle regeneration in the VML model through the utilization of artificial materials. This achievement not only establishes a novel methodology within the field of muscle tissue engineering but also presents novel therapeutic modalities for addressing muscle defects resulting from trauma, surgical interventions, or congenital abnormalities.

## Materials and methods

### 1. Fabrication of collagen hernia patch

#### 1.1. Production of collagen thread

The collagen thread was fabricated using soluble collagen. After dissolving the solid collagen into a solution, it was extruded from a spinneret (e.g., a syringe needle) into a coagulation bath to form collagen threads. Subsequently, these threads were cross-linked. For further details, please refer to an article [He et al., 2025a].

The collagen thread exhibits remarkable mechanical strength and possesses exceptional biological properties. It can function as an absorbable surgical suture, as officially recognized by the Chinese Food and Drug Administration. (CFDA) (No. MZ17010224, No. WT17080974, No. WT19010144, No. WT19010145).

#### 1.2. Production of hernia patch

Collagen threads, approximately 0.15 mm in diameter, are employed for plain weaving. The warp density is set at 140/10 cm, the weft density at 1140/10 cm, and the total warp count is 40. Subsequently, a sheet of 30 × 30 mm is prepared, with a thickness about 0.40 mm. This sheet will be utilized for repairing the VML in the rabbit abdomen.

### 2. Mechanical strength of hernia patches

#### 2.1. Tensile strength

Trim the hernia patch into 5 mm wide strips. Attach two ends of a strip to the mechanical testing machine fixture. Stretch the strip at a uniform speed of 100 ± 10 mm/min until it breaks, and record the maximum force. Soak a 5 mm wide strip in physiological saline for 1 hour and measure its strength as described above.

#### 2.2. Suture strength

5-0 nylon suture is passed through at 5mm at the edge of one end of the hernia patch sheet, and two ends of the nylon suture are secured on the mechanical testing machine fixture. Stretch uniformly at a speed of 100±10mm/min until the nylon thread is pulled out of the torn hernia patch sheet, and record the maximum force. Submerge the hernia patch in physiological saline for an hour, and subsequently measure the suture strength, as previously outlined.

### 3. Animal experiment

Animal procedures were strictly adhered to applicable laws, regulations, and institutional guidelines. Animal welfare and ethics were observed rigorously. Experimental animals were housed in a facility with a 12/12 light cycle, a temperature of 22±1°C, and a humidity of 50%. Food and drink were provided on an on-demand basis. This animal experiment was approved by the Ethics Committee of Wuhan University Central South Hospital (No. 2019007). Experimental animals would be sacrificed by administering an overdose of anesthesia.

#### 3.1. Animals

New Zealand white rabbits, ordinary grade, weight 20∼2.7kg, were from Hubei Experimental Animal Research Center.

#### 3.2. VML model

A combination of 10% hydrated chloraldehyde (1 mL/kg peritoneal injection) and fast-sleep-new II (0.3 mL/kg intramuscularly) is administered as general anesthesia. In the central region of the rabbit’s abdomen, a 3×3 cm segment of the abdominal wall muscle, encompassing the entire layer, is excised. The excised abdominal wall is subsequently washed with physiological saline and preserved for evaluation of tensile strength and suture strength.

#### 3.3. Transplantation

The hernia patch is implanted to repair VML by using 5-0 nylon suture. Subsequently, 5-0 nylon suture is used to sew the fascia, and 3-0 silk suture is used to close the epidermal skin wound. After the surgery, penicillin of 100,000 units is injected intramuscularly twice a day for three consecutive days.

#### 3.4. Post transplantation

In the fourth, eighth, sixteenth, twenty-fourth, and thirty-second weeks following surgery, five rabbits were randomly euthanized using excessive anesthesia. Subsequently, the surgical site was cut open, and the surrounding tissue status, as well as the healing of the abdominal wall anastomosis, was observed. Hernia patches containing surrounding tissue (including regenerated tissue) were then excised and preserved. Four samples were utilized to measure the mechanical strength and degradation rate of the implants, while one sample was used for histological analysis. The tensile strength of the regenerated abdominal wall was measured as previously described.

#### 3.4. Historical analysis

Hematoxylin eosin (HE) stain and Masson stain of regenerated abdominal wall tissue was performed.

#### 3.5. Determination of degradation rate of hernia patch

Cut open the regenerated abdominal wall to extract the residual collagen fibers of the hernia patch. Thoroughly rinse and dry the collagen fibers. Measure its weight and calculate the degradation rate of the hernia patch.

### 4. Statistics

All data were presented as Mean ± SD and analyzed using Microsoft Excel; t-test was utilized to compare the two groups; p < 0.05 indicates a statistically significant difference.

## Result

### Morphology of hernia patch

A comparative analysis of the morphological structure of the collagen hernia patch and the rabbit’s abdominal wall is presented in Figure 1. Figure 1a depicts a photograph of the collagen hernia patch, Figure 1b presents its microscopic image, demonstrating that the hernia patch is composed of numerous collagen threads. Figure 1cd illustrates that a collagen thread is composed of regularly arranged multiple small collagen fibers. Figure 1e depicts the rabbit abdominal wall after formalin immersion, revealing the numerous muscle bundles. Figure 1f further illustrates these separated muscle bundles, Figure 1ef all showcases the densely arranged muscle fibers that comprise the rabbit’s abdominal wall muscles. Figure 1g presents Masson staining of abdominal wall muscle bundle, corroborating the neatly arranged muscle fibers within the bundles. This comparative analysis demonstrates that the collagen hernia patch effectively emulates the intricate structure of abdominal wall muscles.

**Figure 1.**
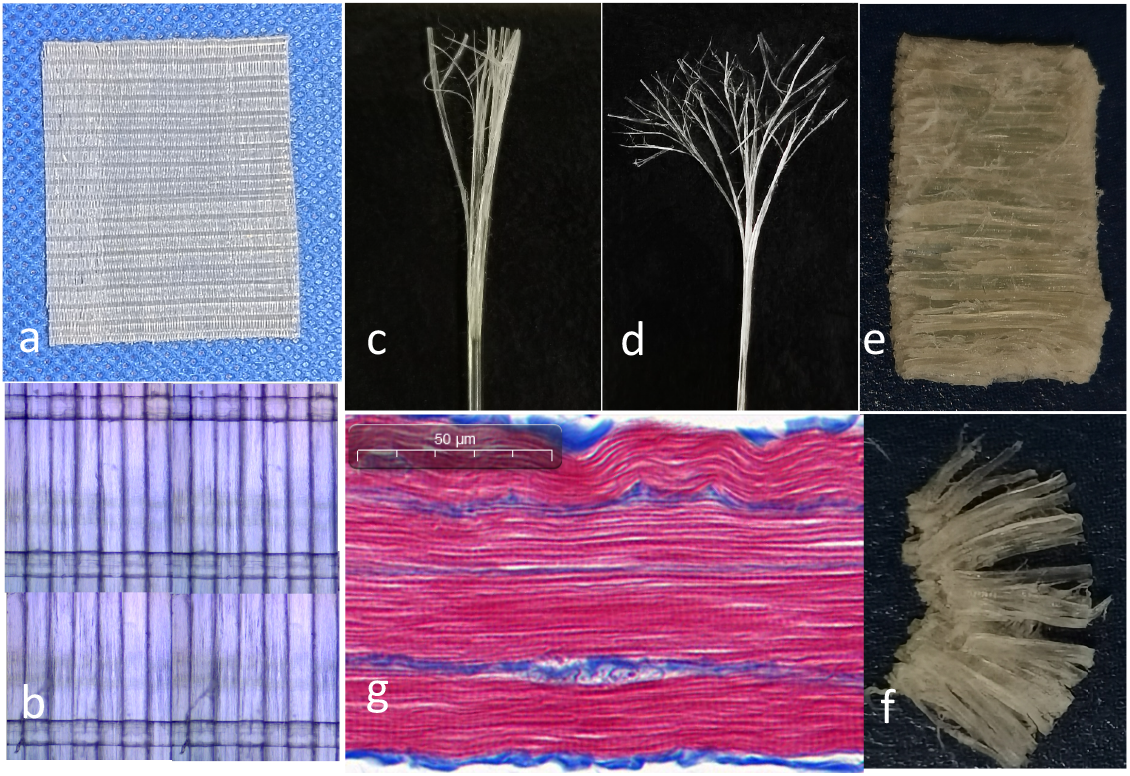
Morphological comparison of collagen hernia patch and abdominal wall muscles. Note: Image c and d were also included in our prior work of tendon regeneration (He et al., 2025b)

### Mechanical strength of hernia patch

In Figure 2, mechanical strength is compared between a collagen hernia patch and a rabbit abdominal wall. The tensile strength (left) and suture strength (right) of the dry patch exhibit a significantly higher value compared to the rabbit abdominal wall. After immersing in physiological saline, the patch’s strength significantly decreases. However, the wet patch retains a non-significantly greater value compared to the rabbit abdominal wall.l.

**Figure 2.**
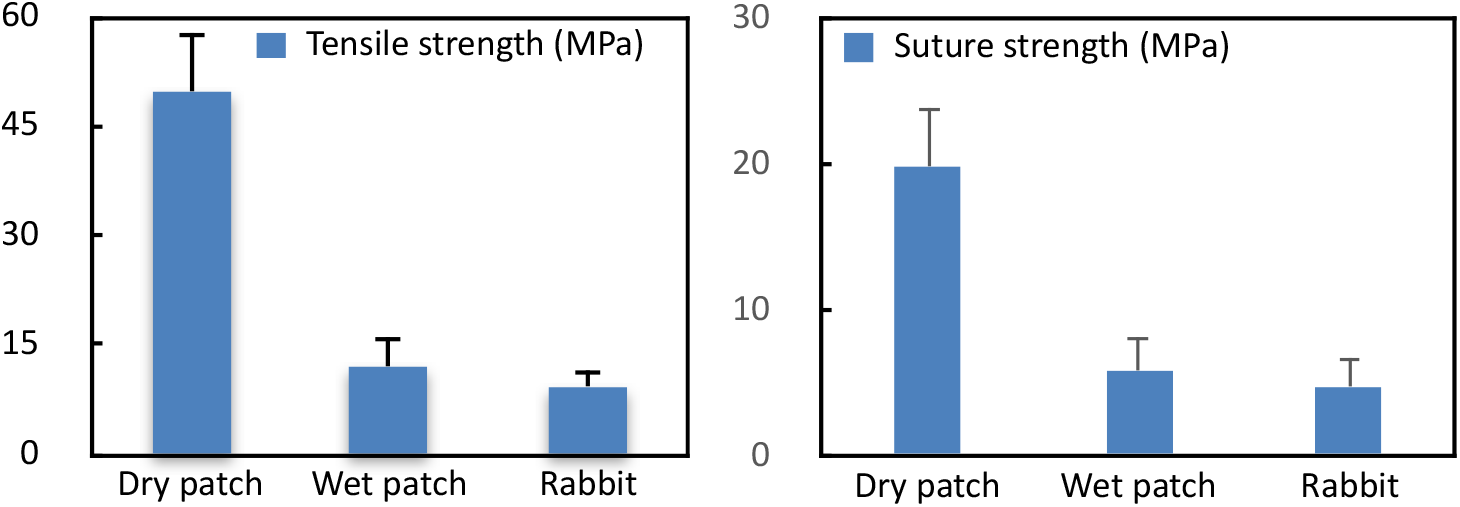
Tensile strength (Left) and suture strength (Right) of the hernia patch and rabbit abdominal wall. Rabbit: Rabbit abdominal wall;

Based on the mechanical strength data of the collagen hernia patch, it is evident that its strength is sufficient to guarantee the safety and reliability of animal experiments. Throughout the entire animal experimental cycle, the implanted hernia patch exhibited no shedding, further demonstrating its stability and safety.

### Transplantation of hernia patch

Figure 3abc illustrates the surgical procedure employed to repair a 3×3 cm VML in the rabbit abdomen using a collagen hernia patch. The surgical site exhibited satisfactory healing within a week, and hair growth subsequently covered the area within three weeks (Figure 3def). The overall appearance was well restored, indicating a favorable wound healing process both superficially and subcutaneously. The surgical site was reopened at 4-week and 8-week post-surgery, and no adhesion to visceral organs was observed (Figure 3gh).

**Figure 3.**
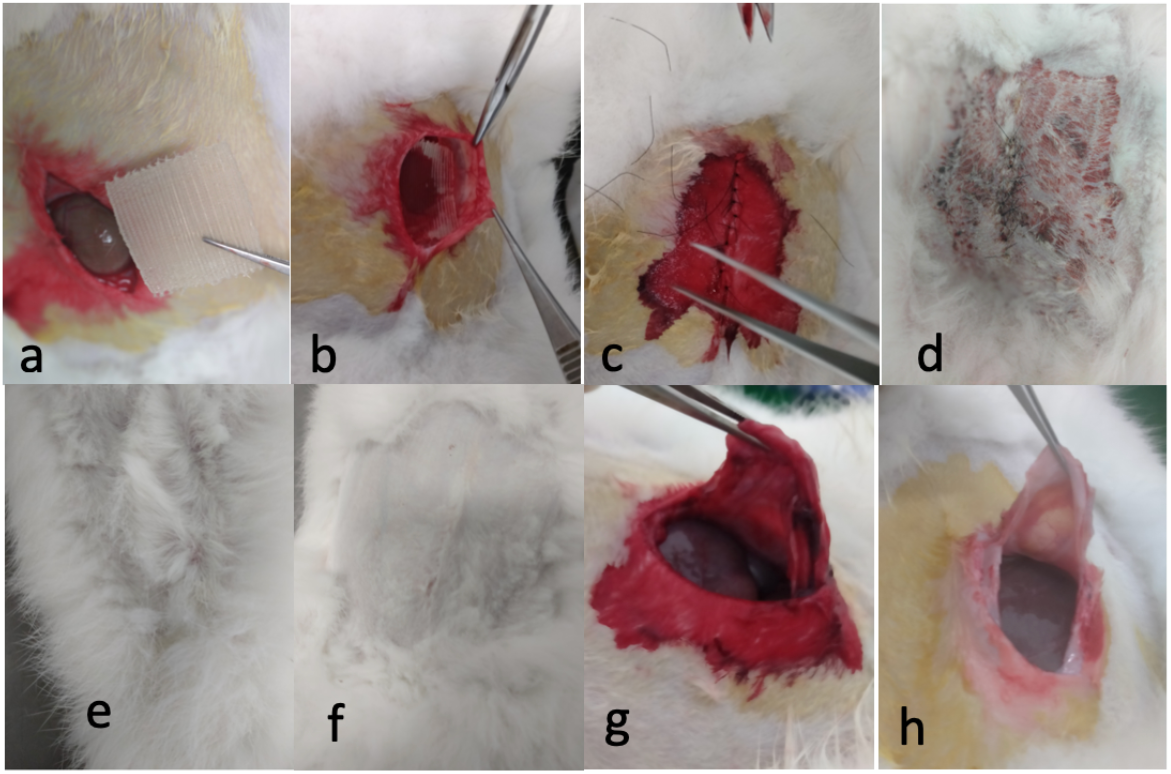
Transplantation process of a collagen hernia patch and the rabbit’s appearance after surgery.

### Collagen hernia patch in vivo

In vivo assessment of the collagen hernia patch was conducted at 17 days post-operation (Figure 4). The epidermis was excised, revealing a newly formed membrane tissue covering the collagen patch (Figure 4a). Subsequently, the covering tissue was meticulously separated from the collagen patch (Figure 4bc), enabling clear visualization of the patch in the eye (Figure 4c). Finally, the collagen patch was fully separated from the underlying newly formed tissue without causing damage to the patch or the natal tissue (Figures 4de). Following the removal of the collagen patch (Figure 4f), new vessels and blood were observed on the underlying tissue (Figure 4e). Blood spots were also observed in the hernia patch (Figure 4f), indicating that the collagen patch recruits capillaries or facilitates the formation of new vessels. Notably, newly formed tissue was regenerated both on and beneath the collagen hernia patch, and can be easily separated from the collagen patch, indicating that the regenerated tissue is not adhering scar tissue. It demonstrates that the collagen hernia patch can recruit functional cells to the defect area and promotes the formation of new vessels, thereby accelerating tissue regeneration and wound healing. The regenerated tissue effectively separates the visceral organ from the hernia patch, eliminating any potential adhesion.

**Figure 4.**
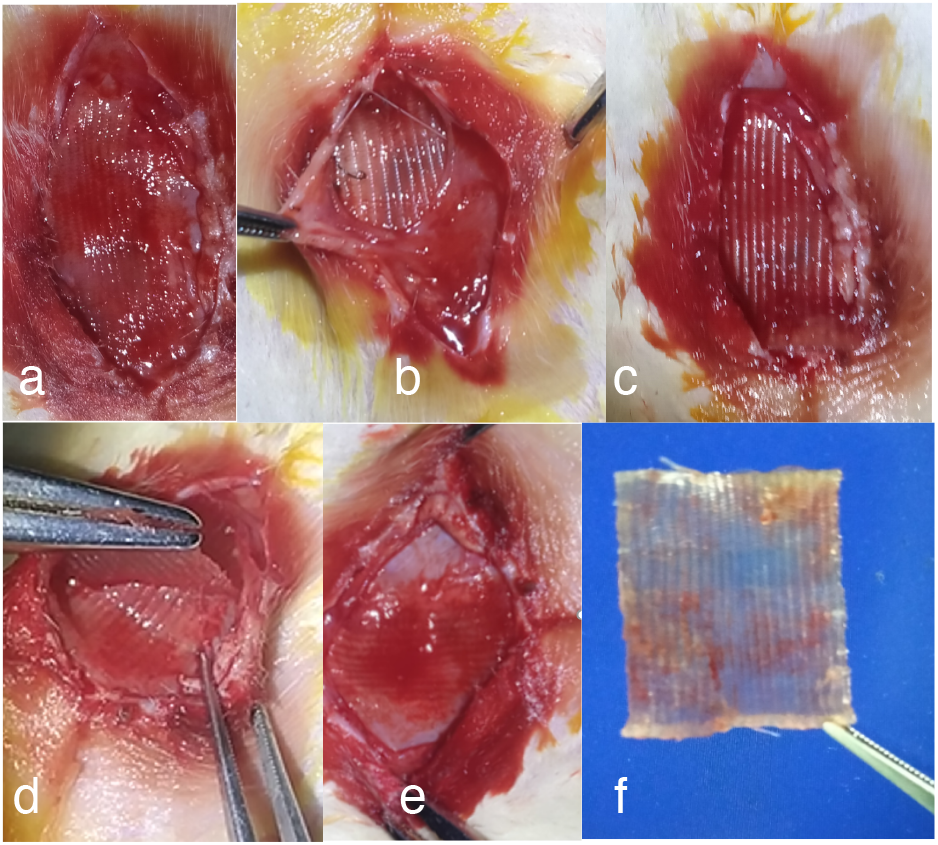
In vivo hernia patch at 17-day postoperative

### Regenerated abdominal wall

Figure 5 (left panel) depicts the external surface, internal surface, and cross sections of the regenerated abdominal wall 16 weeks postoperative, there is no discernible difference from the natural autologous abdominal wall,

**Figure 5.**
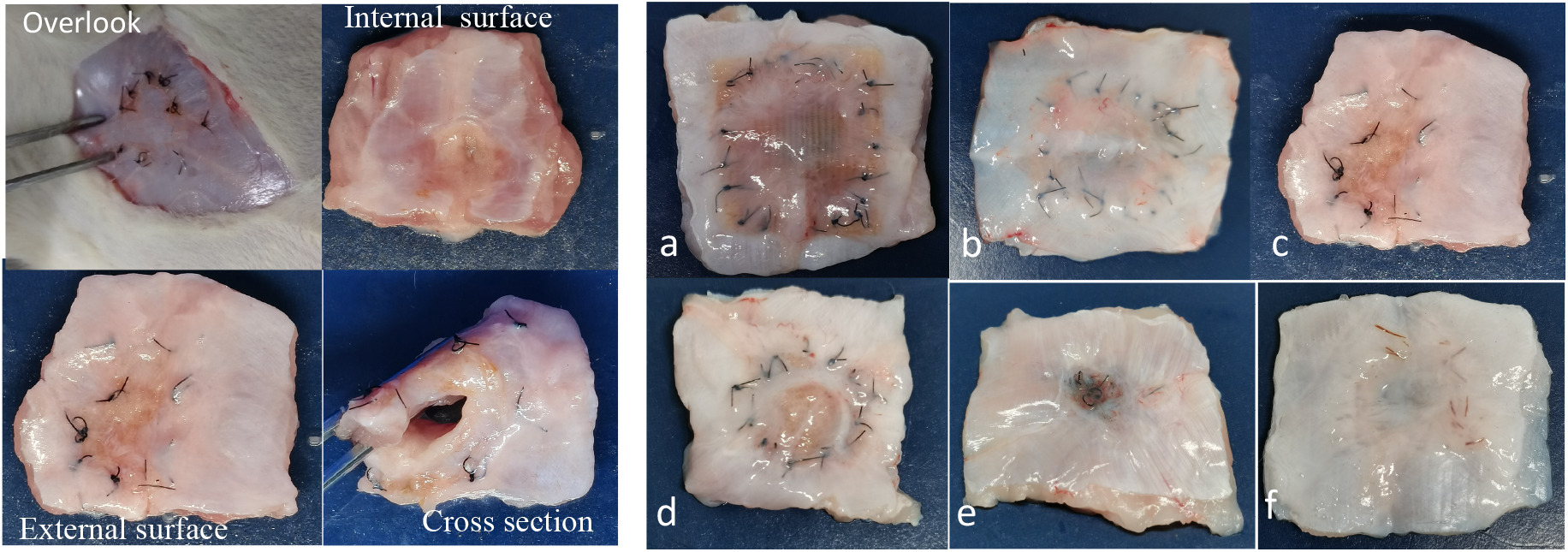
Left panel: a sample of regenerated abdominal wall from 4 perspectives. Right panel: the regenerated abdominal walls in chronological order.

The sequential images of the regenerated abdominal wall tissues (external surface) are presented in Figure 5 (right panel), illustrating the progressive regeneration process. Notably, as time progresses, the nylon sutures gradually relocate from the anastomosis towards the VML center. By approximately 32 weeks post-surgery, they have concentrated to the center, creating a black dot in the VML center (Figure 5e). The collagen hernia patch and its degradation products stimulate the recruitment of stem cells and capillaries to the defect site, facilitating the regeneration of muscle tissue. The regenerated muscle tissues progressively expand inward, propelling the nylon sutures forward. Consequently, the surgical sutures are displaced forward and concentrated at the VML center.

Morphological analysis of the regenerated tissue reveals the absence of a distinct boundary surrounding the anastomosis. Furthermore, scar tissue is not observed around anastomosis, suggesting that the muscle tissue has been successfully regenerated and seamlessly integrated with autologous muscle tissue. Figure 5abcde demonstrates the application of nylon sutures to connect the hernia patch to repair the VML. Black absorbable collagen thread (suture) was also utilized to connect the hernia patch to repair the VML. The surgical suture undergoes degradation over time, resulting in the absence of a black dot (nylon sutures) in the center of the 32-week sample (Figure 5f). However, residual remnants of the black collagen suture remain around the center (Figure 5f). It appears that additional time is necessary for the muscle regeneration process (Figure 5f).

### Histological analysis

Figure 6 (ABCDE) depicts a cross-sectional view of the regenerated abdominal wall in chronological order, with the hernia patch embedded within it. The collagen hernia patch are readily distinguishable particularly at 4 weeks and 8 weeks postoperatively (Figure 6AB). Over time, the hernia patch undergoes a gradual process of degradation and absorption, ultimately disappearing at approximately 24 weeks (Figure 6DE).

**Figure 6.**
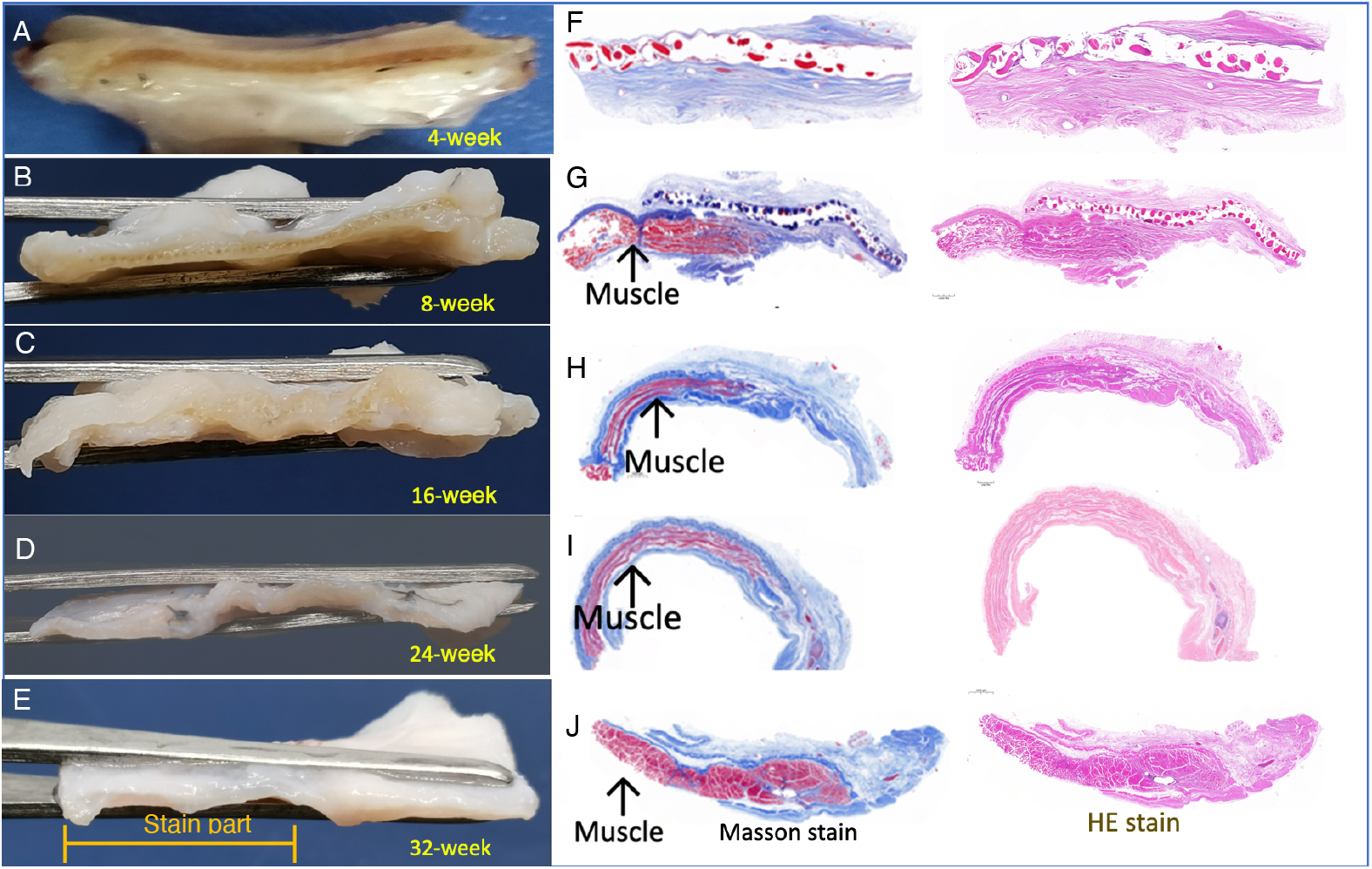
Regenerated abdominal wall in chronological order. Left column: cross sections of abdominal wall; Middle column: Masson stain (blue); Right column: HE stain (pink). Half of the newly formed tissue was used for HE stain and Masson stain, as indicated in image E.

Masson stains muscle tissue red, while other tissue appears blue. Figure 6 (FGHIJ middle column) illustrates a Masson stain of regenerative abdominal wall tissue sequentially. At 4 weeks after collagen hernia patch transplantation, there is still no regenerated muscle tissue surrounding the hernia patch, except for the hernia patch itself being embedded within the tissue (Figure 6F). However, at 8 weeks after transplantation surgery, red regenerated muscle tissue has grown beneath the hernia patch (Figure 6G), and the area of new muscle increased over time (8 weeks → 16 weeks → 24 weeks → 32 weeks) (Figure 6 GHIJ), indicating the regeneration of muscle fibers from the anastomosis to cover the defect. At 32 weeks post-surgery, a substantial mass of muscle tissue is observed, however, muscle tissue is absent at the center of VML (Figure 6J). The concentrated nylon sutures appear to impede muscle regeneration, particularly in the later stages. The collagen content of the hernia patch gradually diminishes over time, yet it remains readily distinguishable during the initial stages, particularly at the 4-week and 8-week after surgery. The hernia patch disappeared approximately 24 weeks after surgery due to its progressive degradation and absorption.

As illustrated in Figure 6 (Right column), the HE stain of the regenerating abdominal wall exhibits a uniform red (pink) coloration. This coloration presents a challenge in distinguishing muscle tissue from other tissues. However, the collagen hernia patch is clearly discernible at both the 4-week and 8-week stages.

### In vivo mechanical strength and degradation rate

Figure 7(left) depicts the alterations in the mechanical strength of the regenerated abdominal wall following transplantation of the hernia patch. The strength exhibited a rapid decline, reaching its lowest point at the 8-week. Subsequently, it gradually rebounded, showing a relatively stable strength increase in the regenerated abdominal wall. Notably, the strength in the 32-week closely resembled that of the autologous abdominal wall.

**Figure 7.**
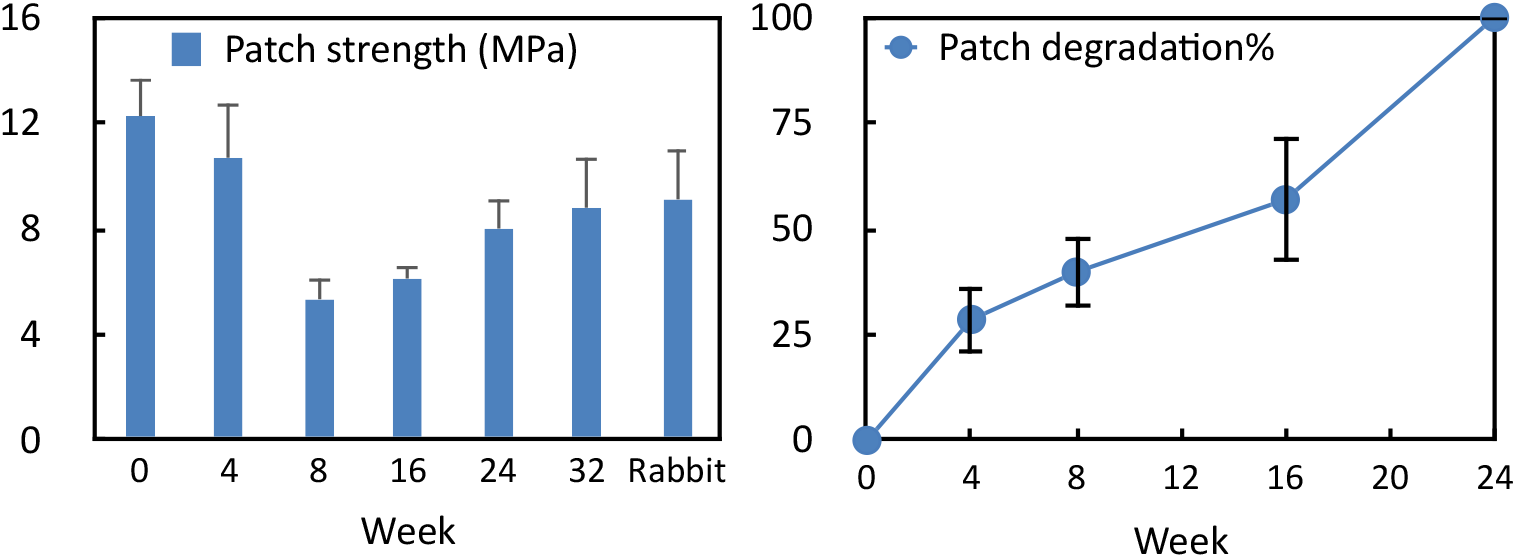
Left: Mechanical strength of regenerated abdominal wall. Rabbit: Rabbit abdominal wall; Right: In vivo degradation rate of the hernia patch.

Figure 7 (right) illustrates the chronological degradation of the hernia patch in vivo. During the initial 16 weeks in vivo, the degradation rate remains relatively low and stable. Subsequently, the degradation rate accelerates, coinciding with the strength rebound at approximately 16 weeks. After 16 weeks, the degradation rate undergoes a significant increase, culminating in complete degradation at 24 weeks postoperative.

## Discussion

Volumetric muscle loss (VML) has long been recognized as one of the most formidable obstacles within the field of tissue engineering.

Satellite cells remain dormant between the fibrous basal membrane and the plasmic membrane of muscle tissue, which primarily contribute to muscle growth and play a crucial role in repairing minor muscle damage[Chargé & Rudnicki, 2004; Yin et al., 2013]. However, the repair ability of satellite cells is limited in VML area, the local microenvironment, such as hypoxia, inflammatory reactions, and insufficient nutrient supply, hinders the migration, proliferation, and differentiation of satellite cells [Sacco et al., 2008; Lepper et al., 2011]. If VML is not adequately treated, a large number of fibroblasts, macrophages, neutrophils, and other immune cells would firstly and rapidly fill the defect, resulting in the formation of scar tissue and the subsequent loss of local function [Jia et al.,2020; Wang et al.,2020].

The number of satellite cells remains relatively fixed and limited, they cannot span the damage gap and cover the entire defect area without a proper scaffold. Muscle regeneration also necessitates a complex network of precisely regulated growth factors, cytokines, and other signaling molecules. Maintaining the network in VML area is challenging, which negatively impacts the function of satellite cells [Sacco et al., 2008; Lepper et al., 2011].

Consequently, it is imperative to utilize excellent biomimetic materials akin to autologous muscle tissue to repair the defects. This present study utilizes high-affinity, low-antigenic soluble collagen to construct a high-strength biological scaffolds as hernia patches. This methodology facilitates muscle regeneration of VML (3×3 cm defect) in the rabbit abdominal wall, thereby achieving a substantial advancement in the field.

### Mechanisms of VML muscle regeneration

VML exceeds the size that the body can heal naturally and is referred to as a “critical defect.”[Testa et al., 2021]. The hernia patch based on soluble collagen achieves functional muscle regeneration in repairing VML, Its intrinsic mechanism can be mainly attributed to the biomimetic properties of collagen hernia patch.

Firstly, the collagen hernia patch, devoid of antigens such as, cellular DNA, α-gal, and collagen telopeptides, can be recognized as “self” by the immune system, thereby significantly reducing inflammatory and immune responses, minimizing the likelihood of scarring or fibrosis [Badylak et al., 2009]. The collagen material induces macrophages to polarize to the M2 phenotype through the TLR4 signaling pathway, releasing anti-inflammatory factors such as TGF-β and IL-10, simultaneously activates Smad and PI3K/Akt pathways to induce stem cells to differentiate into muscle cells [Murray 2017; Zhang et al., 2010]. Collagen has remarkable adsorption capabilities facilitate its synergistic combination with various growth factors, such as integrins (e.g., α2β1, α5β1), IGF-1, HGF, and VEGF, to establish a localized high-concentration bioactive microenvironment, in such microenvironment, collagen scaffolds and its degradation byproducts play a pivotal role in recruiting stem cells to the damaged site through CXCR4/SDF-1 pathway, FAK/MAPK signaling pathway, and can further activate the Notch, Wnt, and Smad signaling pathways, guiding stem cells to migrate, proliferate, and differentiate into muscle fibers [Li et al., 2004; Engler et al., 2006; Guilak et al., 2009; Schwartz & Ginsberg, 2002; Levenberg et al., 2005; Zammit et al., 2004]. In a word, collagen and its byproducts provide a suitable microenvironment for functional cells activities. Martinello et al. and Zheng et al. also demonstrated the regenerative material should be able to promote the proliferation and differentiation of endogenous stem cells/progenitor cells, and supports the concept of “material-guided in vivo tissue engineering”.

In this study, collagen fibers of hernia patch emulates the muscle fiber arrangement, this topological structure can directs stem cells’ expansion, migration, proliferation, and differentiation in a specific direction, resulting in a well-structured muscle tissue [Choi et al., 2008; Angelina et al.,2018; Hochleitner et al., 2018; Yin et al., 2010; Zheng et al., 2017]. Additionally, our advanced cross-linking technology substantially enhances the tensile strength of the hernia patch. This material not only provides mechanical support during the initial phase of cell recruitment but also maintains its strength to facilitate functional muscle reconstruction until collagen material degradation and complete muscle regeneration. Additionally, cross-linking hinders the degradation of collagenase and matrix metalloproteinase (MMP), thereby synchronizing the degradation rate of the hernia patch with tissue remodeling and the regeneration of new tissue [O’Brien, 2011].

After implantation of hernia patch, revascularization and infiltrating of stem cells (satellite cells etc.) initiate within the biomimetic materials, collagen and its degradation products play a crucial role in stimulating the formation of new blood vessels, which is essential for tissue repair and muscle regeneration [Davis and Camarillo, 1995; Ruhrberg et al.,2002]. Additionally, M2 macrophages secretes VEGF, FGF to promote angiogenesis, by inducing endothelial cell migration and form a new capillary (diameter 10–50 µm), ensuring the blood supply of regenerated tissue [Murray, 2017: Levenberg et al., 2005]. Collagen short peptide (<10 kDa) enhances the stability of new blood vessels through the VEGF/ANG-1 signaling pathway and prevents ischemic collapse, which is particularly important for large-volume tissue regeneration [Wang et al., 2009]. Neural integration is the pivotal link to muscle restoration, the structural characteristics of collagen fibers can mimic the mechanical environment of muscles, providing physical cues that support the growth of nerve axons and facilitate the reconstruction of nerve-muscle junctions [Jessen and Mirsky, 2016; Kjaer, 2004; Chen et al., 2007]. Notably, VEGF not only stimulates angiogenesis but also contributes to the release of neurotrophic factors, facilitating the integration of neurons into newly formed muscle tissue [Gonzalez et al., 2011].

### Three phenomena in the research

In the scenario of minor superficial muscle injuries, satellite cells assume a pivotal role in the repair process. functional repair can attain a state of near-perfect regeneration, resulting in minimal scarring[Chargé & Rudnicki, 2004; Yin et al., 2013]. However, in the case of VML, scar formation seems an unavoidable consequence.

In this study, polypropylene (PP) were also employed in the repair of the VML. However, severe adhesion to internal organs was observed in almost each PP case (Figure 8). This is because the damage area of VML is extensive, and the inflammatory response is so severe that scarring process (adhesion formation) is inevitable. As a non-degradable synthetic material, PP can induce inflammatory and immune reactions, leading to the proliferation of fibrous tissue and adhesion to neighboring organs. The rough surface of the PP patch facilitates the adhesion of cells and tissues, thereby increasing the likelihood of adhesion formation. Additionally, implanting a PP hernia patch alters the local microenvironment, disrupts blood flow, and elevates oxidative stress, all of which contribute to the promotion of adhesion formation [Arung et al., 2011]. Severe adhesion necessitates surgical intervention, which, regrettably, may potentially compromise the adhesion. In contrast, under identical surgical conditions, adhesion was not observed when using a collagen hernia patch in this study. There was no discernible immune or inflammatory response.

**Figure 8.**
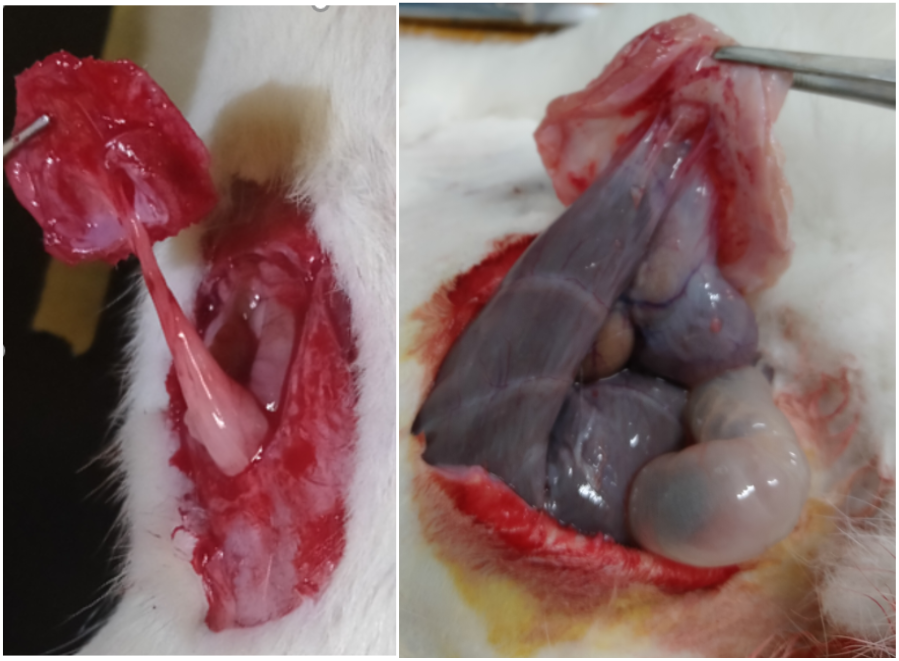
The robust adhesion exhibited by PP in repairing rabbit VML models.

Additionally, two other noteworthy phenomena are also observed in the study. The first one is the migration and concentration of the nylon suture toward the defect center during muscle regeneration, which is primarily attributed to the movement of reconstructed muscle cells, including myocytes [James et al., 2002; Gabbiani et al., 2003]. As mentioned previously, the collagen hernia patch and its degradation products recruit and facilitate functional cells expansion, migration to the defect site, and promote the formation of new blood vessels by creating an ideal microenvironment [Engler et al., 2006; Guilak et al., 2009; Davis and Camarillo, 1995; Ruhrberg et al., 2002]. The migration, proliferation and differentiation of these cells propel the non-degradable nylon sutures forward.

The another phenomenon is illustrated in Figure 5, depicting the regeneration of soft tissue both on and beneath the collagen patch, the regenerated tissues effectively prevent internal organs from adhering to the collagen patch.

All 3 phenomena suggest that the collagen hernia patch exhibits low antigenicity, high compatibility, and high ‘nutritional value’, thereby recruiting many active cells and facilitating functional muscle regeneration rather than tissue scarring. Its composition, function, and structure, which are comparable to natural tissue, ensure its seamless integration into the host body.

Soluble collagen undergoes a pre-treatment process that effectively eliminates telopeptides. Furthermore, our materials technology has ingeniously modified the molecular spatial structure of soluble collagen, thereby further reducing its antigenicity. The hernia patch also aligns collagen fibers in the direction of muscle fibers to construct a mechanically guided spatial structure. All these factors enhance the regenerative potential of satellite cells and inhibit the fibrotic processes of fibroblasts. Consequently, VML healing is facilitated, and scarring is minimized.

Notably, this material transcends its role as a physical support structure, serving as a source of cellular ‘nutrition’. In comparison to synthetic polymers PLA, PGA and others or decellularized matrix, its biological activity exhibits significantly heightened levels.

### Biomimetic structure

One of the important performance of this biomimetic collagen patch is exhibited as a supportive structure for muscular regeneration.

The structural biomimetics in tissue engineering is intriguing. For instance, this study successfully regenerated a 2-3 mm thick abdominal wall using a single-layer 0.4 mm collagen patch. However, there is a significant disparity between 0.4 mm and 2 mm, and the 0.4 mm collagen hernia patch also lacks the delicate structure of natural autologous muscle. This suggests that the fine structure of regenerated muscle fibers was not determined by the collagen hernia patch in this study, it is highly probable that the muscle regeneration process was predominantly guided by the inherent genetic blueprint of satellite cells and the interplay between stem cells and the surrounding microenvironment.

We have also employed a four-layer collagen patch(1.6 mm thick) to repair VML, which is closer to the structure of natural rabbit abdominal wall. Surprisingly, despite using this artificial scaffold to repair VML for enhanced muscle tissue regeneration, the outcome was not as favorable as that achieved with a single-layer 0.4 mm collagen patch. It appears that the four-layer structure is not advantageous for muscle tissue regeneration. The specific reasons for this non-function remain to be investigated.

The conventional method in tissue engineering emulating tissue structure and constructing highly precise artificial scaffolds, overlooks the fundamental biological principles governing stem cell growth and differentiation. Stem cells primarily adhere to the blueprint encoded in cellular DNA (and the local microenvironment) rather than solely relying on the guidance of intricate scaffold structures. Despite the precisely structural emulation attained in scaffold design employing materials like metal and ceramics, these structures lack biological activity for tissue development control [Wang et al.,2020; Hwangbo et al., 2021], functional cells (including stem cells) cannot flourish in an environment containing much metal or ceramics that lacks the optimal conditions for cell migration, growth, and differentiation. Consequently, their regenerative potential remains largely unexplored. Experiments on this muscle regeneration demonstrate that even on a rudimentary collagen scaffold (0.4 mm thick hernia patch), stem cells can spontaneously form highly intricate functional muscle tissues, offering novel concepts for the simplified design of regenerative medical materials.

Furthermore, this study has demonstrated the presence of numerous small vascular tissues surrounding regenerated muscles, which also originate from the internal body system rather than from the structure of the transplanted hernia patch.

Consequently, providing an exceptional biomimetic material that closely resembles autologous tissue is the optimal choice. The biomimetic collagen patch successfully emulates the composition, structure, and function of the abdominal wall, thereby creating an environment conducive to stem cell development, i.e., this biomimetic scaffold offers a comprehensive biological, mechanical, nutritional, and other favorable microenvironment that facilitates the migration and differentiation of satellite cells (stem cells) into muscle fibers in situ. This process enables the repair of VML by the formation of new muscle tissue [Jia et al.,2020; Wang et al., 2020].

This study emphasizes the strategy of “activating endogenous self-repairing capabilities” over excessive reliance on simulating organizational structures. The fundamental principle of this study aligns with the developmental biology concept that each cell possesses a comprehensive genome “design blueprint.” Provided with a suitable microenvironment, cells will naturally differentiate into their designated tissue type.

### Research limitations

Although the findings of this study are promising, there are several critical concerns that warrant our immediate attention. (1)Individual Variation: Most rabbits exhibit successful muscle regeneration, while some exhibit slow or even incomplete regeneration. These disparities may be attributed to individual health conditions, genetic background, and surgical procedures. To address this, we should implement further standardization of materials and techniques to minimize subjective and man-made variables. (2) Suture Material Impact: The impact of nylon sutures on the muscle regeneration process is substantial. Nylon sutures occupy the space for muscle regeneration and may induce local inflammation, which can impede the regeneration process. Consequently, surgical sutures should be comprehensively evaluated within the broader context of the regeneration system. (3)Large Animal Verification: Although the preliminary results obtained in the rabbit model are promising, their validity has not been substantiated in large animal models such as pigs and dogs. These latter models are more physiologically similar to humans in terms of abdominal wall muscle volume, stress load, and immune environment, making them crucial for clinical translation. (4) Human Application: The ultimate objective is to utilize the technology for clinical practice in human applications. Initially, the biomimetic materials can be employed to repair minor muscle defects, such as hernias.

### Future research

It is important to note that the current research primarily focuses on histological repair effects. In the future, it is imperative to systematically assess: (1) Functional Recovery Assessment: Evaluate the capabilities of regenerated muscles through comprehensive functional tests, including electromyograms, muscle strength assessments, and contractility tests. This assessment will evaluate the overall functional recovery of the regenerated muscles. It will ensure the contraction and diastolic properties of the regenerated skeletal muscles, thereby ensuring their compliance with the desired standards. (2) Muscle Regeneration in Other Regions: Although abdominal wall muscle regeneration has demonstrated efficacy, it is imperative and prudent to investigate muscle regeneration in other regions of the body. The human body comprises 639 skeletal muscles, with those in the palms, arms, and legs being particularly susceptible to damage. The functional regeneration of these muscles holds substantial clinical value and can be initially evaluated in large animal models. (3) Scaffold Material Optimization: Enhance the regeneration ability in ischemic environments by optimizing the structure of the scaffold material, including adjusting the space area, fiber density, and incorporating angiogenic factors. (4) Combined Cell Therapy Exploration: For patients with low immunity or advanced age and infirmity, the joint use of scaffolds, autologous stem cells, inducing factors, and other therapeutic modalities may be considered to potentially enhance the success rate.

Neuromuscular diseases, including Duchenne muscular dystrophy (DMD), amyotrophic lateral sclerosis (ALS), and spinal muscular atrophy (SMA), present intricate etiologies and pose significant therapeutic challenges. In addition to the muscle regeneration observed in this study, this materials technology has demonstrated remarkable regeneration of functional tissue in the major defect of the sciatic nerve and tendon in rabbit models. Consequently, this newly developed materials technology holds substantial medical implications in the treatment of these muscle atrophy diseases by synergizing with advanced technologies such as CRISPR-Cas9, AAV gene therapy, and stem cell technology. One approach involves cultivating stem cells [e.g., satellite cells, neural stem cells(NSCs), neural progenitor cells(NPCs)] within this biomimetic materials and differentiating them into muscle tissues or neuromuscular tissue in vitro or in vivo. Subsequently, this differentiated tissue can be transplanted into human bodies.

### Clinical prospects

The collagen hernia patch scaffold exhibits a simple structure, remarkable biological properties, and has successfully regenerated functional muscle tissues without the need for exogenous cells, growth factors, or immunosuppressants. These attributes facilitate standardization and large-scale production, thereby enhancing its clinical translational potential. Consequently, it holds the promise of widespread application in the following areas: (1) Abdominal wall defect repair (e.g., post-tumor surgery, traumatic injuries); (2) Muscle defects resulting from war injuries or severe traffic accidents;(3) Congenital muscle loss or muscle atrophy lesion reconstruction; (4) Soft tissue reconstruction auxiliary materials in plastic surgery and sports medicine;

Given its accessibility as a readily available medical material, its low cost, and broad clinical applicability, it is anticipated to transform into a groundbreaking product in the field of regenerative medicine.

### Conclusion

This study demonstrates that high-strength biological scaffolds constructed from soluble collagen can effectively achieve VML-level muscle regeneration without the necessity of exogenous cells or intricate biomimetic structures. This finding underscores that the primary factor in regeneration lies not in “replicating tissue structure,” but rather in “activating the potential of stem cells and establishing a favorable microenvironment.” Despite remaining challenges in clinical applications, this study proposes a novel paradigm for advancing next-generation muscle regeneration materials, thereby opening promising avenues for the field of tissue engineering.

